# Interleukin 7 receptor drives Early T lineage Progenitor expansion

**DOI:** 10.1101/2021.10.23.465581

**Authors:** Rafael A. Paiva, Vera C. Martins

**Affiliations:** Lymphocyte Development and Leukemogenesis Laboratory, Instituto Gulbenkian de Ciência, Calouste Gulbenkian Foundation, Oeiras, 2780-156, Portugal

**Keywords:** Early T lineage progenitor, ETP, T lymphocytes, T lymphocyte development, Thymopoiesis, Thymus, Interleukin 7, Interleukin 7 receptor

## Abstract

Interleukin 7 (IL-7) and IL-7 receptor (IL-7r) are essential for T lymphocyte differentiation, by driving proliferation and survival of specific developmental stages. While early T lineage progenitors (ETP), the most immature thymocyte population known, have a history of IL-7r expression, it is unclear whether IL-7r is required at this stage. Here, we show that mice lacking IL-7 or IL-7r have a marked loss of ETPs that results mostly from a cell-autonomous defect in proliferation and survival, although no changes were detected in Bcl2 protein levels. Further, a fraction of ETPs responded to IL-7 stimulation *ex vivo* by phosphorylating Stat5, and IL-7r was enriched in the most immature Flt3^+^Ccr9^+^ ETPs. Consistently, IL-7 promoted the expansion of Flt3^+^ but not Flt3-ETPs on OP9-DLL4 cocultures, without affecting differentiation at either stage. Taken together, our data show that IL-7/IL-7r is necessary following thymus seeding, by promoting proliferation and survival of the most immature thymocytes.

**Summary:** Paiva et al. show that IL-7/IL-7r signaling upon thymus seeding is essential for proliferation and survival of the most immature early T lineage progenitors (ETP), thereby determining the physiological ETP cellularity.

## Introduction

T lymphocytes differentiate from hematopoietic progenitors of bone marrow origin that seed the thymus (Yui and Rothenberg, 2014). The estimated number of thymus seeding progenitors in adult mice is in the range of 150-200, with only ∼10 open niches available for seeding at steady-state (Zietara et al., 2015). The few thymus seeding progenitors proliferate mostly as result of Notch1 signalling and give rise to the most immature thymocyte population, the early T lineage progenitor (ETP) (Hosokawa and Rothenberg, 2021). The ETP is a heterogeneous population composed of sequential subsets that follow a highly dynamic process of gene regulation towards T lineage commitment (Zhou et al., 2019). Within the ETP, the most immature cells express Chemokine Receptor 9 (Ccr9) and Fms Related Receptor Tyrosine Kinase 3 (Flt3) (Benz and Bleul, 2005; Sambandam et al., 2005). These receptors are lost in few days, together with alternative cell fate potentials, as result of Notch1/Deltex4 signalling that instructs T cell lineage commitment (Benz et al., 2008; Feyerabend et al., 2009; Heinzel et al., 2007). Further differentiation of the ETP into the CD4-CD8-double negative (DN) 2 (DN2) stage completes commitment towards the T cell lineage (Koch et al., 2008; Radtke et al., 1999).

Mice deficient for interleukin 7 (IL-7) or either of the IL-7 receptor (IL-7r) chains (IL-7rα, or common gamma chain, γ_c_) cannot generate T lymphocytes (Cao et al., 1995; DiSanto et al., 1995; Peschon et al., 1994; von Freeden-Jeffry et al., 1995). In immunodeficient patients with loss-of-function mutations in the homologous genes T lymphocytes also fail to differentiate (Noguchi et al., 1993; Puel et al., 1998). This reflects the absolute requirement of IL-7/IL-7r signaling for T lymphocyte differentiation (Hong et al., 2012). While IL-7/IL-7r signaling in the adult thymus is best described in DN2 and DN3 thymocytes (Balciunaite et al., 2005; Boudil et al., 2015; Tussiwand et al., 2011), its role at former stages has yet to be addressed. Lineage tracing studies *in vivo* have shown that the ETP has a history of IL-7rα expression (Schlenner et al., 2010). While the label in the ETP was considered to report recombination in upstream bone marrow progenitors, it is difficult to completely exclude IL-7r expression by ETPs (Schlenner et al., 2010). In this line, *IL-7r* expression in the newborn thymus has been reported in 50-55 % of Flt3-positive ETP (Luc et al., 2012). These data led us to ask whether the ETP requires IL-7/IL-7r signaling.

We show that, in the adult thymus, the adult ETP population requires IL-7/IL-7r signaling. Mice lacking IL-7r have a marked loss of ETPs that is mostly caused by defective proliferation and survival. A fraction of ETPs phosphorylated Stat5 *ex vivo* in response to IL-7 stimulation. Further, IL-7r expression was enriched in the most immature ETPs identified by Flt3 and Ccr9. Consistently, IL-7 promoted the expansion of Flt3^+^ but not Flt3-ETPs on OP9-DLL4 co-cultures, without affecting differentiation at either stage. Altogether, our data show that IL-7/IL-7r is required following thymus seeding to promote expansion and survival of the most immature thymocytes, thereby contributing to establish the normal cell number of ETPs in the adult thymus.

## Results and Discussion

### IL-7/IL-7r deficiency causes a marked reduction in ETPs

We noted that the ETP population is greatly reduced in *IL-7rα-/-* adult thymi when compared to wild type controls (Fig. 1A). While a reduction in the absolute number of ETPs in thymi deficient for IL-7 or IL-7r (*IL-7rα-/-* or *γ-/-*) could be due to general thymic atrophy (Fig. 1B), the reduction in frequency (Fig. 1C) hints at a specific mechanism. To test whether the phenotype could be caused by an effect prior to thymus seeding, we quantified the bone marrow progenitors proposed to contribute to thymopoiesis, i.e. common lymphoid progenitors (CLPs) and lymphoid-primed multipotent progenitors (LMPPs). Both progenitor populations were increased in percentage in IL-7 deficient bone marrow relative to wild type controls (Fig. S1A, S1B), which reflects the absence of other cells requiring IL-7/IL-7r signaling, such as mature lymphocytes and B cell precursors. However, the absolute cell number of both CLPs and LMPPs did not differ between mutant and control mice (Fig. S1C, S1D).

**Figure 1.**
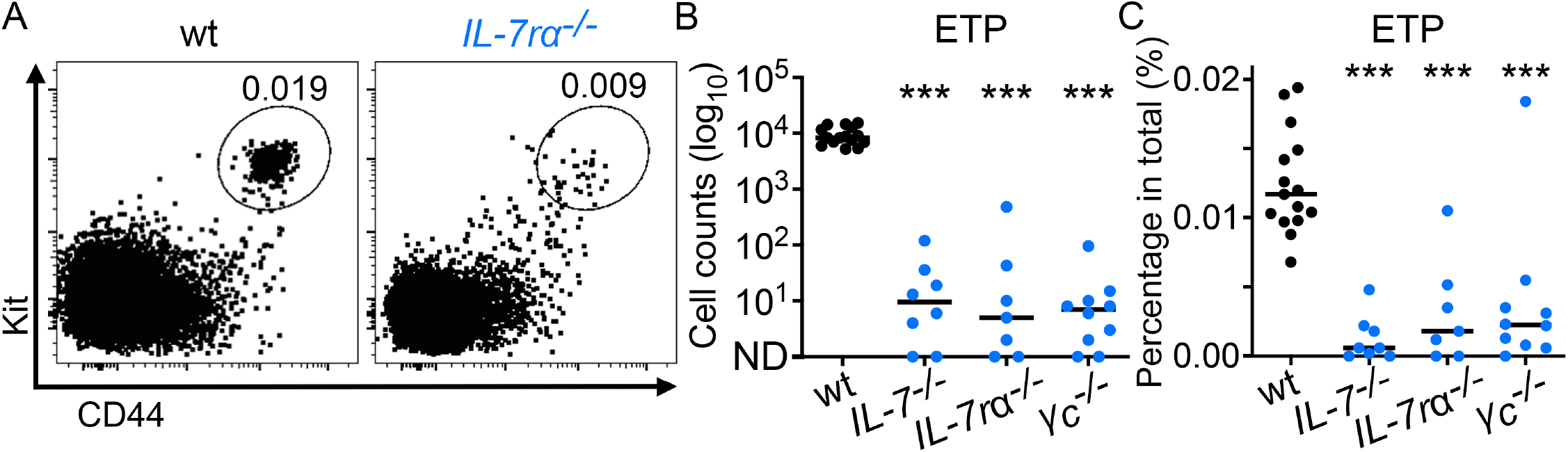
IL-7/IL-7r deficiency results in the loss of ETP. **A)** Wild type and *IL-7rα*^*-/-*^ thymocytes were pre-gated as lineage^-^CD25^-^. The gate identifies the ETP and the numbers are the percentage of ETP in total thymocytes. **B)** Cell counts of ETP in the indicated genotypes, and C) the corresponding percentage of ETP in total thymocytes in 6-week-old mice. Each circle corresponds to one thymus and the lines are the medians. Data are from two independent experiments for each tested genotype and wild type mice were used in every experiment. Statistical significance was calculated with Mann Whitney test: ***p≤0.001. See also Figure S1.

### The ETP cellularity depends mostly on intrathymic IL-7/IL-7r signaling

The identity of the physiological bone marrow progenitor population(s) contributing directly to thymopoiesis remains unresolved (Bhandoola et al., 2007; Krueger, 2018), and it was therefore possible that the bone marrow analyses (Fig. S1) might have missed a minor, but relevant, subpopulation. To test whether the loss of ETPs was caused by a defect prior to thymus homing, we transplanted wild type thymi into either wild type or *IL-7-/-* recipients (Fig. 2A). In these experiments, progenitors developed either in the presence or absence of IL-7 and were exposed to IL-7 only upon thymus seeding. In both conditions, cells are IL-7r proficient and can, therefore, respond to IL-7 after thymus colonization that is produced by the wild type thymus stroma. Thymus grafts were analyzed 5 weeks later, ensuring complete repopulation by bone marrow-derived (host) cells. A sizable ETP population was detected in both types of recipients (Fig. 2B). While the total cellularity of the thymus grafts did not differ (Fig. S2A), there was on average a 1.5-fold reduction in percentage (Fig. 2B) and absolute number (Fig. 2C) of ETPs when the host was deficient for IL-7. Importantly, no differences in proliferation (Fig. S2B) or cell death (Fig. S2C) were noted in ETPs regardless of the host genotype. These data suggest that there is a minor role for IL-7/IL-7r signaling in pre-thymic progenitors that affect ETP cellularity. Nevertheless, the pre-thymic defect uncovered with these thymus transplantation experiments could not account for the magnitude of the phenotype observed in IL-7 and IL-7r deficient thymi (Fig. 1A-C). Therefore, to test for a putative effect of IL-7/IL-7r signaling in the ETP solely after thymus seeding, we grafted wild type and *IL-7-/-* thymi into wild type recipients (Fig. 2D). In these conditions, pre-thymic progenitors developed in an IL-7 proficient environment and the effect of IL-7 signaling on ETPs could be revealed following thymus graft seeding. Interestingly, despite having a continuous supply of wild type bone marrow progenitors, the ETP population was almost absent in the *IL-7-/-* thymus grafts in percentage (Fig. 2E) and absolute cell numbers (Fig. 2F). As expected, thymocyte cellularity in the *IL-7-/-* grafts was much lower when compared to wild type thymus grafts (Fig. S2D). Because differences in thymus cellularity could impact on the effective number of niches available for colonization by hematopoietic progenitors, we generated competitive bone marrow chimeras at a 1:1 ratio, whereby wild type progenitors could reconstitute the thymus to its normal size, and *IL-7rα-/-* versus control ETP could be compared (Fig. 2G). As expected, thymus cellularity was similar between the two groups (Fig. 2H). Likewise, no differences were noted in bone marrow reconstitution, with similar engraftment for both wild type and *IL-7rα-/-* progenitors (Fig. S2E). While the disadvantage of *IL-7rα-/-* thymocytes relative to the wild type counterparts was expected, the fact that it started in ETP supported the hypothesis that the population directly depends on *IL-7rα* (Fig. S1H). Indeed, the size of the ETP population was similar in the two experimental groups, with a difference only in the relative contribution of *IL-7rα* proficient or deficient ETPs (Fig. 2I). Altogether, our data indicate that the reduction of ETPs in IL-7/IL-7r deficient thymi results mostly from an intrathymic, cell-autonomous defect.

**Figure 2.**
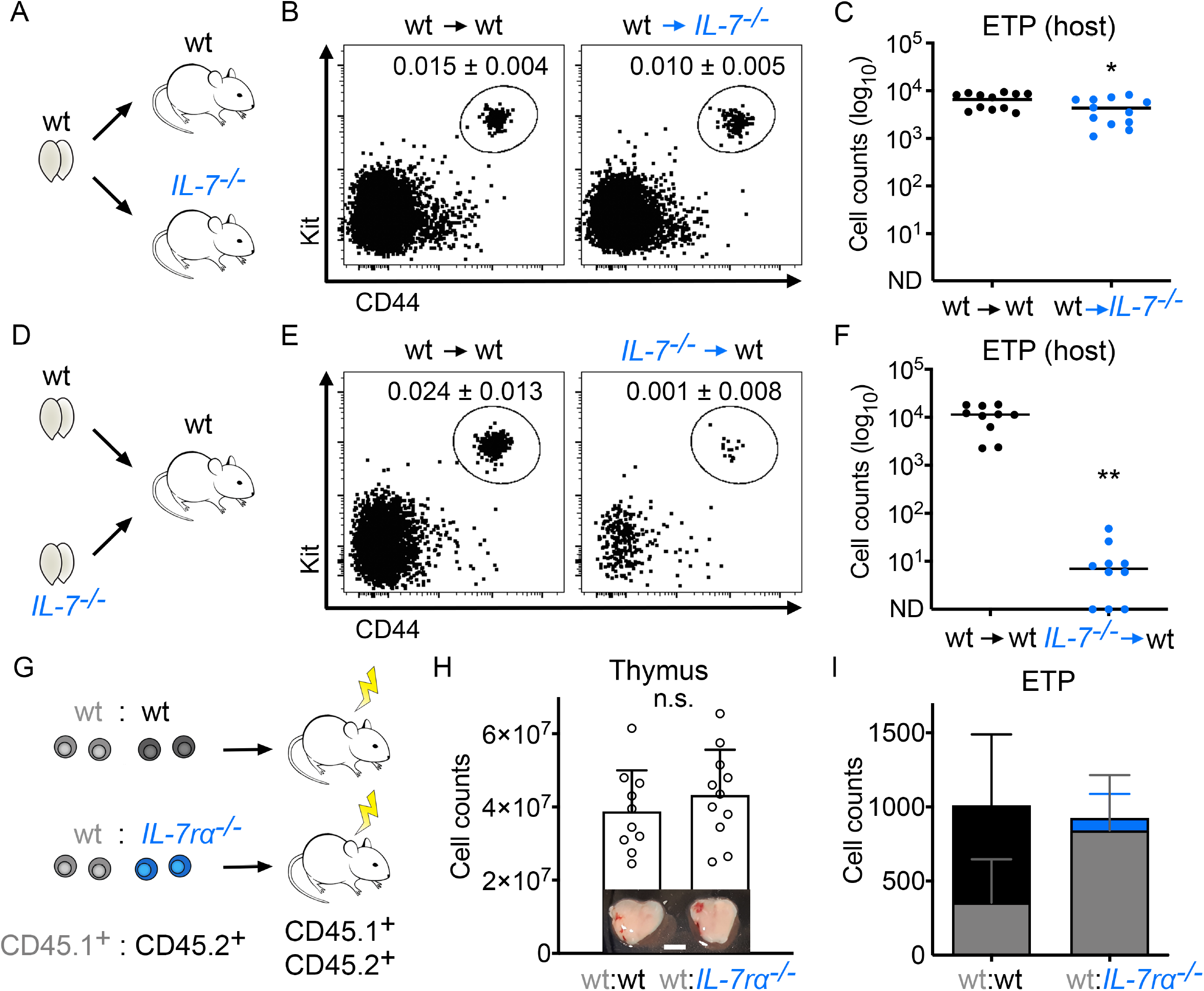
Lack of IL-7/IL-7r cause a cell-autonomous defect in ETPs. **A)** Wild type thymi were transplanted into wild type or *IL-7*^*-/-*^ mice and analyzed 5 weeks thereafter. **B)** Representative FACS plots showing host-derived lineage-CD25-thymocytes, and numbers indicate the percentage (mean ± SD) of ETPs in host-derived thymocytes. **C)** Total number of host-derived ETP in the indicated conditions. **D)** Wild type or *IL-7*^*-/-*^thymi were transplanted into wild type mice and analyzed 5 weeks thereafter. Each recipient received two grafts, one from each genotype. **E)** Representative plots showing host-derived CD4-CD8-lineage-CD25-thymocytes, and numbers indicate the percentage (mean ± SD) of ETP in total host-derived thymocytes. **F)** Cellularity of host-derived ETP in wild type (black) or *IL-7*^*-/-*^ (blue) thymi grafted in wild type recipients. Each circle corresponds to one thymus graft (F, I) and the lines indicate the medians. **G)** Lethally-irradiated wild type mice (CD45.1^+^CD45.2^+^) were reconstituted with wild type bone marrow progenitors mixed at a 1:1 proportion with either wild type or *IL-7rα*^*-/-*^ cells and analyzed 9-10 weeks thereafter. **H)** Total thymus cellularity of the reconstituted mice. Photographs depict one representative thymus for each group. Scale bar = 2.5 mm. **I)** ETP cellularity in thymi from the wild type:wild type chimeras (CD45.1^+^ wild type cells in grey and CD45.2^+^ wild type cells in black) or the wild type:*IL-7rα*^*-/-*^ chimeras (CD45.1^+^ wild type cells in grey and CD45.2^+^ *IL-7rα*^*-/-*^ cells in blue). Data from two independent experiments are depicted as mean ± standard deviation. Statistical significance was calculated with Mann Whitney (C), Wilcoxon signed rank (F) or T test (H): n.s.>0.05, *p≤0.05, **p≤0.01. See also Figure S2.

### IL-7/IL-7r signaling induces proliferation and promotes survival of ETPs, independently of Bcl-2

IL-7/IL-7r signaling in the thymus is known to drive proliferation and survival of DN2 and DN3 thymocytes. Thus, using the DN2 as an internal control, we asked whether these parameters differed between wild type and *IL-7rα-/-* ETPs. *IL-7rα-/-* ETPs proliferated less than their wild type counterparts, as measured by Ki-67 staining (Fig. 3A, B). Additionally, the *IL-7rα-/-* ETP was enriched for dying cells, as determined by the levels of cleaved caspase 3 (Fig. 3C, D). These results were in accordance with those measured in DN2 thymocytes (Fig. 3A-D). However, while IL-7r-deficient DN2 expressed lower levels of Bcl-2, no differences were detected in Bcl-2 protein level between wild type and *IL-7rα-/-* ETPs (Fig. 3E, F). Finally, we tested whether wild type ETPs are responsive to IL-7, and whether signal transduction could be mediated by Stat5. Indeed, we found that a fraction of ETPs phosphorylated Stat5 in response to IL-7 (Fig. 3G, H). While this differed from the homogeneous response of the DN2, it shows that a subset of ETPs is responsive to IL-7. Altogether, these data show that IL-7/IL-7r signaling in ETP promotes proliferation and survival, although the latter is not mediated by Bcl-2 expression.

**Figure 3.**
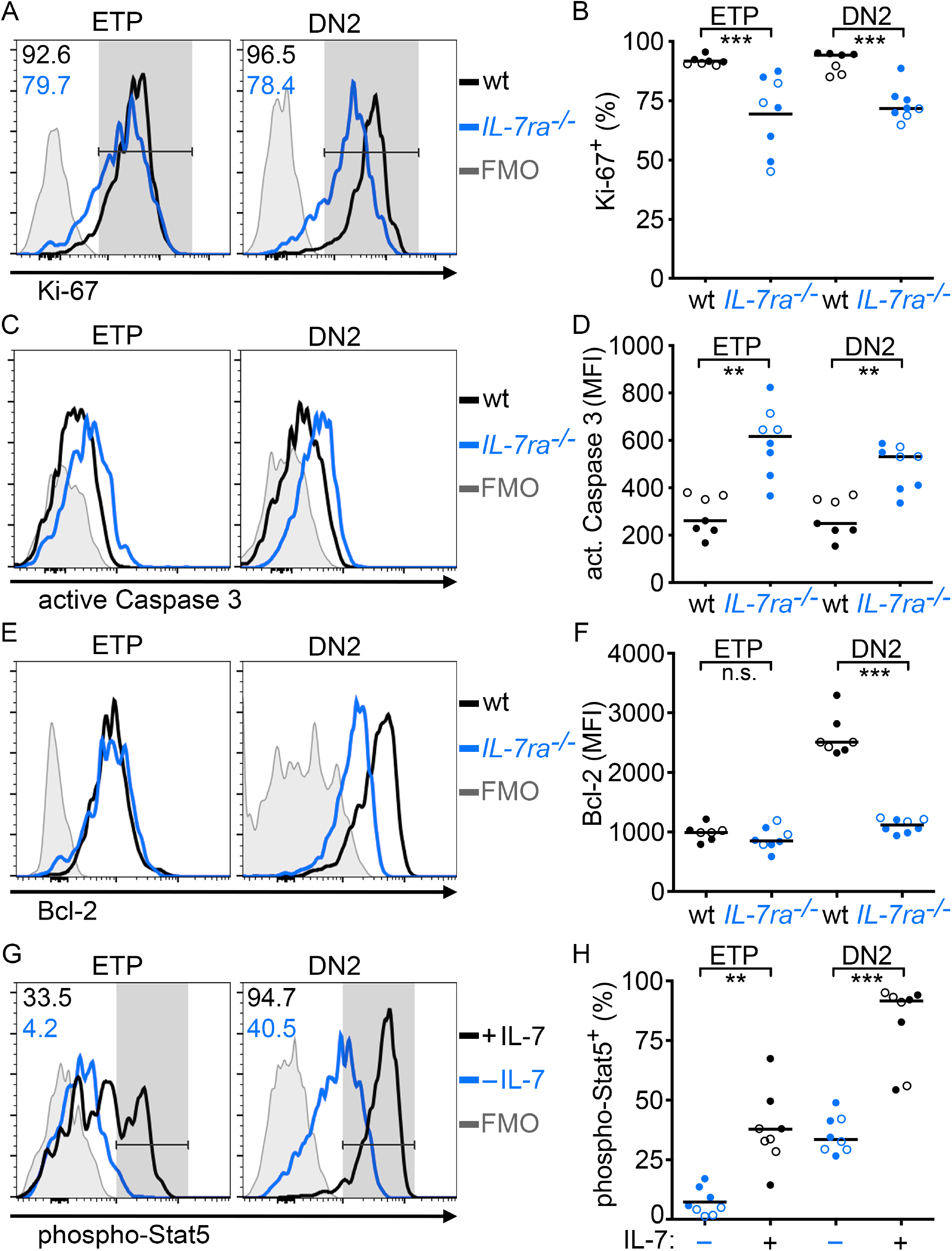
IL-7r promotes proliferation and survival of the ETP. **A)** Ki-67 expression in ETP and DN2 thymocytes from wild type (black) or *IL-7rα*^*-/-*^ (blue) mice. **B)** Percentage of Ki-67^+^ wild type (black) or *IL-7rα*^*-/-*^ (blue) cells in the indicated populations. **C)** Histograms of active caspase 3 in ETP and DN2 thymocytes from wild type (black) or *IL-7rα*^*-/-*^ (blue) mice. **D)** Mean fluorescence intensity of active caspase 3 in the wild type (black) or *IL-7rα*^*-/-*^ (blue) thymocyte populations indicated. **E)** Bcl-2 expression in ETP and DN2 thymocytes from wild type (black) or *IL-7rα*^*-/-*^ (blue) mice. **F)** Mean fluorescence intensity of Bcl-2 in the wild type (black) or *IL-7rα*^*-/-*^ (blue) thymocyte populations indicated. Lines indicate the medians and each symbol represents one thymus for wild type and pools of 2 to 4 thymi for *IL-7rα*^*-/-*^. **G-H)** Wild type thymocytes were cultured for 15 minutes with or without IL-7 and Stat5 phosphorylation was analyzed. **G)** Stat5 phosphorylation in ETP or DN2 without (blue) or following IL-7 stimulation (black). **H)** Percentage of phospho-Stat5^+^ ETP or DN2 after culture without (blue) or with IL-7 (black). Each circle (across conditions) corresponds to one mouse thymus and the lines indicate the means. FMO controls are shown in grey and data are from two independent experiments discriminated by the full or open symbols. Statistical significance was calculated with Mann Whitney (B, D, F) and paired t (H) tests: ^n.s.^p>0.05, **p≤0.01, ***p≤0.001.

### IL-7/IL-7r signaling induces proliferation of the most immature ETPs without affecting differentiation

Since only a fraction of ETPs responded to IL-7, we reasoned that receptor expression might be heterogeneous and addressed whether this could be related to differentiation. Within the ETP, we used Flt3 expression to discriminate the most immature cells (Sambandam et al., 2005) (Fig. 4A), and found that they were enriched in IL-7r-expressing cells (Fig. 4B, C). IL-7r expression in the ETP was lower than in DN2, and did not correlate with the differentiation from ETP to DN2a (Fig. S3A, B). Indeed, IL-7rα expression was downregulated from Flt3hi to Flt3-ETP, and was only upregulated as thymocytes reached the DN2a stage (Fig. S3A, B). These data were consistent with bulk RNAseq data generated by others (Zhou et al., 2019), and reanalyzed here, that showed a similar expression pattern for *IL-7r* mRNA (Fig. S3C). We further assessed heterogeneity within the ETP with basis on Ccr9 protein expression and reporter expression using Ccr9*eGFP* mice (Benz and Bleul, 2005) (Fig. 4D) and consistently found that IL-7r expression is enriched in the most immature ETPs (Fig. 4E, Fig. S3D). Next, as Flt3-positive ETPs were larger than Flt3-negative (Fig. 4A), we asked whether this reflected differences in proliferation. Indeed, proliferation was highest in Flt3hi ETPs and became progressively lower as ETPs downregulated Flt3 (Fig. 4F, G). Of note, Flt3-positive ETPs were virtually absent from *IL-7rα*^-/-^ thymi (Fig. 4H, I). This suggests that Flt3-positive ETPs are more sensitive to the loss of IL-7/IL-7r. To test this hypothesis, we sorted Flt3-positive versus Flt3-negative ETPs (Fig. S3E) and co-cultured them on OP9-DLL4 stromal cells in the presence or absence of IL-7. Flt3-positive ETPs generated CD25 positive cells (transitional to DN2) slower than Flt3-negative (Fig. 4J, Fig S3F), as expected from their respective developmental stages. Nevertheless, the differentiation kinetics from either Flt3-positive and Flt3-negative ETPs was independent of the presence of IL-7 in the cultures (Fig. 4J, Fig. S3F). Instead, IL-7 promoted ETP expansion, with a more pronounced effect on Flt3-positive ETPs, rather than the more mature, Flt3-negative ETPs (Fig. 4K). Altogether, our data indicate that IL-7/IL-7r signaling promotes expansion of the most immature ETPs while not affecting their differentiation.

**Figure 4.**
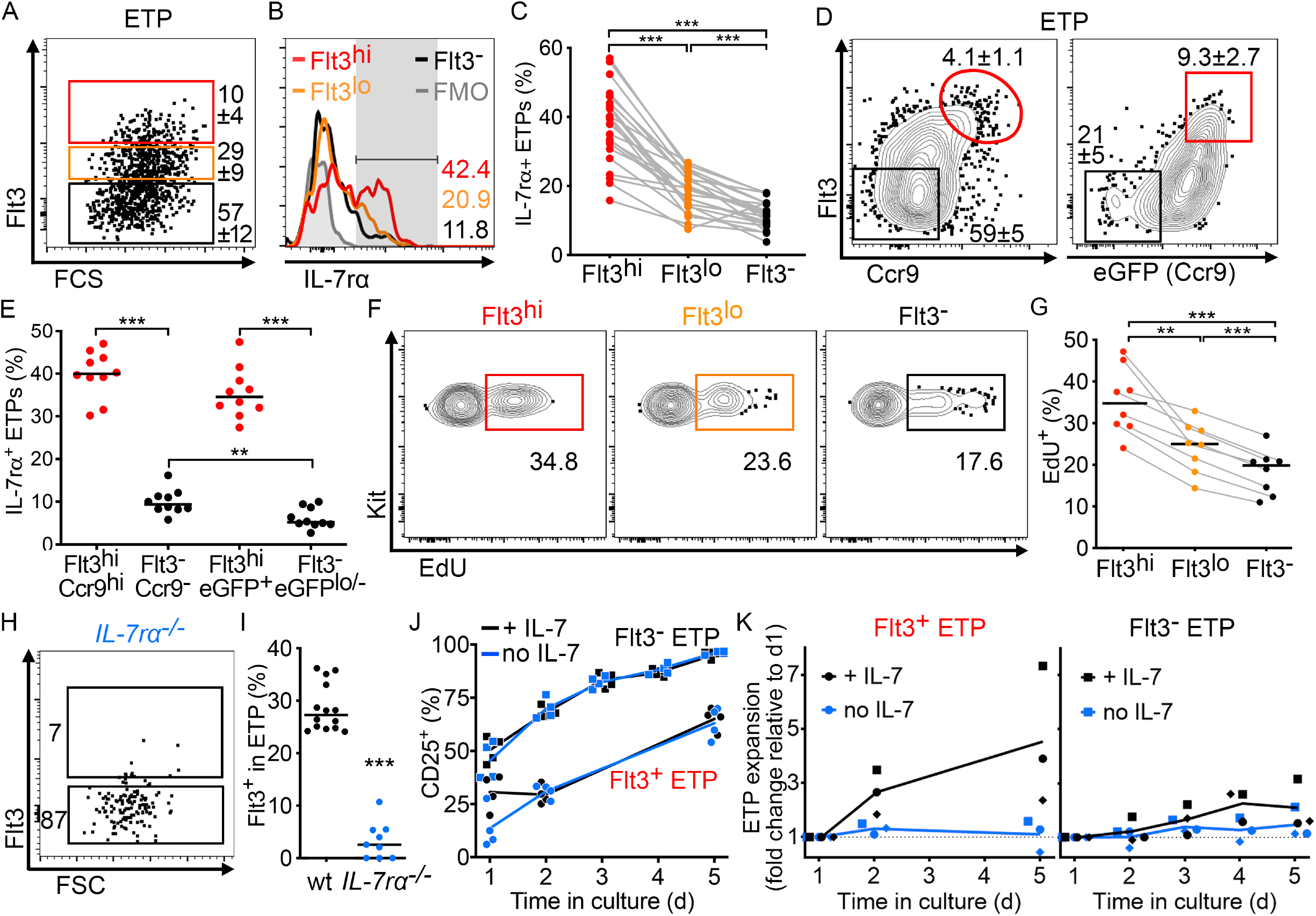
The most immature Flt3^+^ ETP proliferates more at steady-state and expands *ex vivo* in response to IL-7. **A)** Three to 5-week-old thymi were analyzed and shown are the gates used to define Flt3hi, Flt3lo, and Flt3-ETP. Numbers indicate the percentage (mean ± SD) of ETPs. **B)** Subpopulations in (A) were analyzed for IL-7rα expression and an FMO control is shown in grey. **C)** Percentage of IL-7rα^+^ cells in the indicated populations. Data from seven independent experiments (n=22); grey lines connect the subpopulations from individual thymi. C) Thymi from 3 to 5-weeks-old wild type (left) or Ccr9eGFP (right) mice were analyzed and shown are contour plots for Flt3 *versus* Ccr9 (antibody staining or eGFP reporter expression, respectively) pre-gated on ETP. Numbers indicate the percentage (mean ± SD) of ETPs. **E)** Percentage of IL-7rα^+^ cells in the indicated ETP subpopulations. Data are from 3 independent experiments. Each dot corresponds to one thymus and the lines mark the medians. **F)** Three to 4-week-old wild type mice were injected i.p. twice with EdU 2 hours apart and analyzed 2 hours after the last injection. EdU incorporation is shown in the ETP subpopulations. **G)** Percentage of EdU^+^ cells in the indicated ETP subpopulations. Data from 3 independent experiments, the grey lines connect the subpopulations from individual thymi, and the black lines represent the mean. **H)** ETP from 3-to 5-weeks-old *IL-7rα*^*-/-*^ mice were analyzed for Flt3 expression, and **I)** the percentage of Flt3-positive ETP was quantified in wild type (black) or *IL-7rα-/-* (blue) mice. Each symbol represents one thymus for wild type, pools of 2-5 *IL-7rα*^*-/-*^ thymi, and the lines indicate the medians. Data from 3 independent experiments. **J)** Flt3^+^ or Flt3^-^ETP were co-cultured on OP9-DLL4 and the percentage of CD25^+^ cells that differentiated in the cultures was quantified over time in the presence (black) or absence of IL-7 (blue). Data corresponds to the experiment shown in Figure S3B and each dot represents one replicate well. **K)** The same experiments in (J) were used to quantify the expansion of ETP (CD25^+^ cells excluded) with (black) or without IL-7 (blue). The average number of ETP in culture at day 1 was used to normalize the data from the replicates. Data are from 3 independent experiments, each identified by a different symbol, and the lines connect the means. Statistical significance was calculated with RM one-way ANOVA with Tukey’s multiple comparisons (C, G), paired T (E) or unpaired T test (E, Flt3^-^Ccr9^-^ vs Flt3^-^eGFPlo^/-^) and Mann Whitney (I) tests: **p≤0.01, ***p≤0.001. See also Figure S3.

In summary, we show that IL-7/IL-7r signaling is an essential and non-redundant driver of survival and proliferation at the ETP stage, in particular for the most immature Flt3-positive progenitors. We propose that the rare progenitors seeding the thymus use IL-7/IL-7r together with Notch1/Deltex4 signaling to survive and expand, thereby contributing to the generation of ETPs in physiological numbers. We show that genetic abrogation of either the cytokine or the receptor results in a dramatic reduction in the number of ETPs, which is in line with previous reports (Jensen et al., 2008; Sitnicka et al., 2007). The phenotype resulted mostly from an intrathymic and cell-autonomous defect, with only a minute pre-thymic contribution. Whether IL-7/IL-7r signaling in the bone marrow could instruct progenitors towards the T cell lineage and/or favor their migration to the thymus remains to be tested. Along this line, Notch1 signaling has been proposed to prime bone marrow progenitors towards the T lymphocyte lineage (Chen et al., 2019).

The reduced number of ETPs in IL-7r deficient thymi might explain why these thymi are permissive to direct colonization by hematopoietic progenitors injected into the bloodstream (Prockop and Petrie, 2004). While classical experiments hinted at a regulation of thymus seeding by competition for free niches, it is plausible that the niches are unoccupied in IL-7r deficient thymi because ETPs fail to expand. In this context, IL-7/IL-7r signaling might be explored as a potential means to improve thymus function following bone marrow transplantation. In patients undergoing bone marrow transplantation, infections, relapse, and graft versus host disease are common complications that associate with the long recovery time of the T lymphocyte compartment (Castermans et al., 2011; Hakim et al., 2005). The thymus has therefore received considerable attention as to how it could be treated to reduce the refractory period of thymopoiesis after bone marrow transplantation (De Barros et al., 2013). While administration of IL-7 in bone marrow transplanted patients led mostly to an expansion of effector memory T cells (Perales et al., 2012), studies in mice show that thymus activity is improved in similar settings (Bolotin et al., 1996; Mackall et al., 2001). It will therefore be important to test whether thymus function can be improved specifically, and whether the regulation of thymus seeding by IL-7/IL-7r signaling improves thymus reconstitution and function following bone marrow transplantation in patients.

## Materials and Methods

### Mice

C57BL/6J (referred to as wild type, CD45.2^+^) were maintained and bred at Instituto Gulbenkian de Ciência (IGC) in a colony frequently refreshed by mice imported from Charles River. B6.SJL-Ptprc3a (B6.SJL) mice were purchased from The Jackson Laboratory (stock no. 002014). *IL-7rα*^-/-^ (stock no. 002295) and *γc*^-/-^ (stock no. 003174) mice were purchased from The Jackson Laboratory. *IL-7-/-* (von Freeden-Jeffry et al., 1995) were backcrossed to B6 (Carvalho et al., 2001) and kept at IGC. *Ccr7*^*-/-*^ *Ccr9*^*-/-*^ were kindly provided by T. Boehm after crossing the original single mutants (Calderon and Boehm, 2011), and kept at IGC. The original *Ccr7*^*-/-*^ (Ma et al., 1998) had been purchased from The Jackson Laboratory, and the *Ccr9*^*-/-*^ (Benz et al., 2004) were generated by targeting the *Ccr9* endogenous locus with *eGFP* generating a *knock-in/knock-out* at the MPI Freiburg, Germany. *Ccr7*^*-/-*^ *Ccr9*^*-/-*^ were crossed to C57BL/6J to obtain *Ccr7*^*+/-*^ *Ccr9*^*eGFP/+*^ mice (here referred to as *Ccr9*^*eGFP*^). Both males and females were used aged 3 to 6-week-old as indicated, and were age and sex matched in each experiment. The mice were kept in individually ventilated cages under SPF conditions. All animal experiments were approved by the Ethics Committee of the IGC – Fundação Calouste Gulbenkian and the Direção Geral de Alimentação e Veterinária (DGAV) and followed the Portuguese and Europeans laws for animal experimentation.

### Thymus transplants

Thymus transplants were performed as described previously (Martins et al., 2014; Martins et al., 2012; Paiva et al., 2021; Ramos et al., 2020). Briefly, thymi were harvested from newborn mice, the lobes separated and each lobe transplanted into one extremity of the kidney, under the capsule. Mice were kept anesthetized with Xylazine (16 mg/kg) and Ketamine (100 mg/kg). Recipient mice were 5- to 8-week-old and groups were age and sex matched in each experiment.

For the experiment comparing wild type and *IL-7-/-* mice grafted with wild type thymi, donors were CD45.1^+^CD45.2^+^ or CD45.1^+^ and recipients were CD45.2^+^. For the experiments comparing wild type and *IL-7-/-* thymus donors, each wild type recipient received one wild type and one *IL-7*^*-/-*^ thymus graft. Wild type donors were CD45.1+CD45.2^+^, *IL-7*^*-/-*^ donors were CD45.1^+^ or CD45.2^+^ and recipients were CD45.2^+^ or CD45.1^+^, respectively.

### Competitive bone marrow chimeras

Bone marrow cells from 5-6 week-old B6.SJL (CD45.1^+^) and wild type or *IL-7rα-/-* (both CD45.2^+^) were depleted of lineage-positive cells by magnetic separation using Biotin Binder Dynabeads (Thermo Fisher Scientific) after staining with biotinylated antibodies against CD3, CD4, CD8, CD11b, CD11c, CD19, Gr-1 and Ter119. Lineage-depleted CD45.1+ cells were mixed with competitor CD45.2^+^ cells at a 1:1 ratio and a total of 1×106 cells per recipient were administered intravenously via tail vein. Recipient mice were wild type (CD45.1^+^CD45.2^+^) with 5-6 weeks of age and irradiated (700 rad) 10-12 hours before reconstitution. Chimeric mice were analyzed 9-10 weeks after reconstitution.

### Flow cytometry analysis and sorting

Organs were harvested in PBS with 10% FBS and cell suspensions were prepared from thymus, by gently smashing against and passing through a 40 μm cell strainer, and from bone marrow, by flushing and disaggregating the marrow with a 26 g syringe before passing through a 40 μm cell strainer. Cells were blocked with mouse IgG (Jackson laboratories) before staining with antibodies listed below as target antigen (clone; fluorophore): Ccr9 (CW-12; AF647), CD3 (145-2C11; APC-Cy7), CD4 (GK1.1; AF647, BV421, BV605, FITC, PE or PE-Cy7), CD8 (53-6.7; APC/Fire750, BV711,FITC or PerCP-Cy5.5), CD25 (PC61; BV421, BV605, FITC or AF594), CD44 (IM7; BV711, eFluor450, FITC or PerCP-Cy5.5), Kit (2B8; APC or APC-Cy7), Flt3 (A2F10; PE), IL-7r (A7R34; PE-Cy7), Sca-1 (D7; PercP-Cy5.5), CD45.1 (A20; APC, BV421 or PE-Cy7), and CD45.2 (104; BV711 or Pacific Blue), all from BioLegend. The lineage cocktail contained CD3 (145-2C11), CD4 (GK1.1), CD8 (53-6.7), CD11b (M1/70), CD11c (N418), CD19 (6D5), Gr-1 (RB6-8C5), NK1.1 (PK136) and Ter119 (TER-119), all conjugated to biotin or PE and from BioLegend. Of note, CD4 was omitted from the lineage when analyzing ETP. Streptavidin conjugated to BV785 or APC-Cy7 (BioLegend) was used to detect biotinylated antibodies. Dead cells were excluded by Sytox Blue (Invitrogen) or Zombie Aqua (BioLegend). Intracellular staining to Ki-67 (16A8; a700) from BioLegend, Bcl-2 (3F11; PE) and active Caspase 3 (C92-605; FITC), both from BD, was performed with True-Nuclear Transcription Factor Buffer Set (BioLegend). Samples were acquired in a BD LSRFortessa X-20 analyzer and all populations analyzed in this study were identified as indicated in Table S1. Sorts were performed in a BD FACSAria II and sorted populations were: Flt3-ETP (Lin^-^CD25^-^CD44^+^KithiFlt3^-^) and Flt3^+^ ETP (Lin^-^CD25^-^CD44^+^KithiFlt3^+^). Populations identified in the manuscript were defined as follows: LSK hematopoietic progenitors (Lin^-^Sca1^+^Kit+IL^-^7r^-^), LMPP (Lin-Sca1^+^Kit+IL^-^7r^-^Kithi), CLP (Lin^-^Flt3^+^IL^-^7r^+^), ETP (CD4^-^CD8^-^Lin-CD25^-C^D44^+^Kithi), DN2a (CD4^-^CD8^-^Lin^-^CD25^+^CD44^+^Kithi), DN2b (CD4^-^CD8^-^Lin^-^CD25^+^CD44^+^Kitlo), DN3 (CD4^-^CD8^-^Lin^-^CD25^+^CD44^-^), DN4 (CD4^-^CD8^-^Lin^-^CD25^-^CD44^-^), CD8 immature single positive (ISP; Lin^-^CD4^-^CD8^+^), CD4 CD8 double positive (DP; CD4^+^CD8^+^), CD4 single positive T cell (SP4; CD3^+^CD4^+^CD8^-^) and CD8 single positive T cell (SP8; CD3^+^CD4^-^CD8^+^).

### Stat5 phosphorylation

Thymi from 4-week-old wild type mice were harvested and 5 million thymocytes per condition were cultured in IMDM medium (Gibco) with 2% FBS (HyClone Thermo Fischer Scientific) for 15 minutes with or without murine IL-7 (50 ng/mL, PeproTech). Following stimuli, cells were immediately fixed with pre-warmed fixation buffer (BioLegend) for 15 min at 37 ºC, permeabilized with chilled True-Phos Perm buffer (BioLegend) for 1 hour at −20 ºC and finally stained for phosphorylated Stat-5 (BD biosciences, clone 47/Stat5; a647) in PBS/10% FBS for 30 min at room temperature. For each thymus 3 samples were prepared: treated with IL-7 but not stained for phospho-Stat5 (FMO control), cultured without IL-7 and stained for phospho-Stat5 and treated with IL-7 and stained for phospho-Stat5.

### EdU incorporation

Three- to 4-week-old mice were injected intraperitoneally twice with 0.25 mg 5-ethynyl-2’-deoxyuridine (EdU; Sigma-Aldrich) 2 hours apart, and thymi were harvested 4 hours after the first injection. Thymocytes were stained for extracellular antigens before EdU detection with click-iT Plus EdU Alexa Fluor 488 Flow Cytometry Assay Kit (Invitrogen), following the manufacturer’s protocol.

### OP9-DLL4 co-cultures

OP9-DLL4 cells were maintained in IMDM medium (Gibco) supplemented with 10% FBS (HyClone), 0.03 % primatone (Sigma), 5 μg/ml insulin (Sigma) and 0.05 mM β-mercaptoethanol (Gibco) before co-culture with thymocytes. On the evening before the sort, OP9-DLL4 cells were irradiated (3000 rad) and 5000 cells/well were plated on 96-well plates. Thymocytes from 3- to 4-week-old wild type mice were stained with biotinylated antibodies for lineage markers (excluding CD4) and depleted of lineage-positive cells by magnetic separation using Biotin Binder Dynabeads (Thermo Fisher Scientific). Flt3+ or Flt3-ETP were pre-sorted for yield and then 200 cells/well were sort-purified directly onto the OP9-DLL4 culture plate. The co-cultures were maintained in IMDM medium supplemented as above but with 2% FBS, murine Flt3L (5 ng/mL) and with or without murine IL-7 (10 ng/mL), both from PeproTech. Cytokines were refreshed every other day.

### Statistical analysis

D’Agostino & Pearson normality test was applied to each dataset. Mann-Whitney, Wilcoxon signed rank, repeated-measure (RM) one-way ANOVA with Tukey’s multiple comparisons, and paired and unpaired T and paired T tests were used, as indicated in the figure legends. Data normality and statistical significance was calculated with Prism 7 and is represented in the figures as follows: ***p≤0.001, **p≤0.01, *p≤0.05, ^n.s.^p>0.05.

## Supplemental Material

Supplemental material are 3 Supplemental Figures and figure legends.

Figure S1. IL-7-deficient mice have normal numbers of lymphoid-biased progenitors.

Figure S2. IL-7r deficiency causes a cell-autonomous defect in ETPs

Figure S3. Dynamics of IL-7rα expression in early T lymphocyte development.

## Author Contributions

RAP designed the project, designed and performed experiments, analyzed data and wrote the manuscript. VCM designed the project, designed experiments and wrote the manuscript.

## Acknowledgements

This work was supported by the Instituto Gulbenkian de Ciência (IGC), Calouste Gulbenkian Foundation, and the Portuguese National Research Council (Fundação para a Ciência e Tecnologia [FCT] Grant PTDC/BIA-BID/30925/2017 to VCM). VCM is supported by an individual contract awarded by FCT (CEECIND/03106/2018). RAP is a PhD student of the IGC Integrative Biology and Biomedicine (IBB) PhD Program and is supported by an individual FCT PhD Fellowship ref. PD/BD/114341/2016. This work had the support of the research infrastructures Congento LISBOA-01-0145-FEDER-022170 and PPBI-POCI-01-0145-FEDER-022122, both co-financed by FCT and Lisboa2020, under PORTUGAL2020 agreement (European Regional Development Fund). The *Ccr7*^*-/-*^ *Ccr9*^*-/-*^ mice that were crossed to obtain the *Ccr9eGFP*, generated by (Benz and Bleul, 2005), were a kind gift from T Boehm (see Methods for details). We thank A Cumano for the OP9-DLL4, and HJ Fehling, J Demengeot and T Boehm for critical reading of the manuscript. We thank the Animal House Facility and the Flow Cytometry Unit of IGC in supporting this work. The authors declare no competing financial interests.

