## Supplemental Figures 1-3 for "Interleukin 7 receptor drives Early T lineage Progenitor expansion"

#### Figure S1

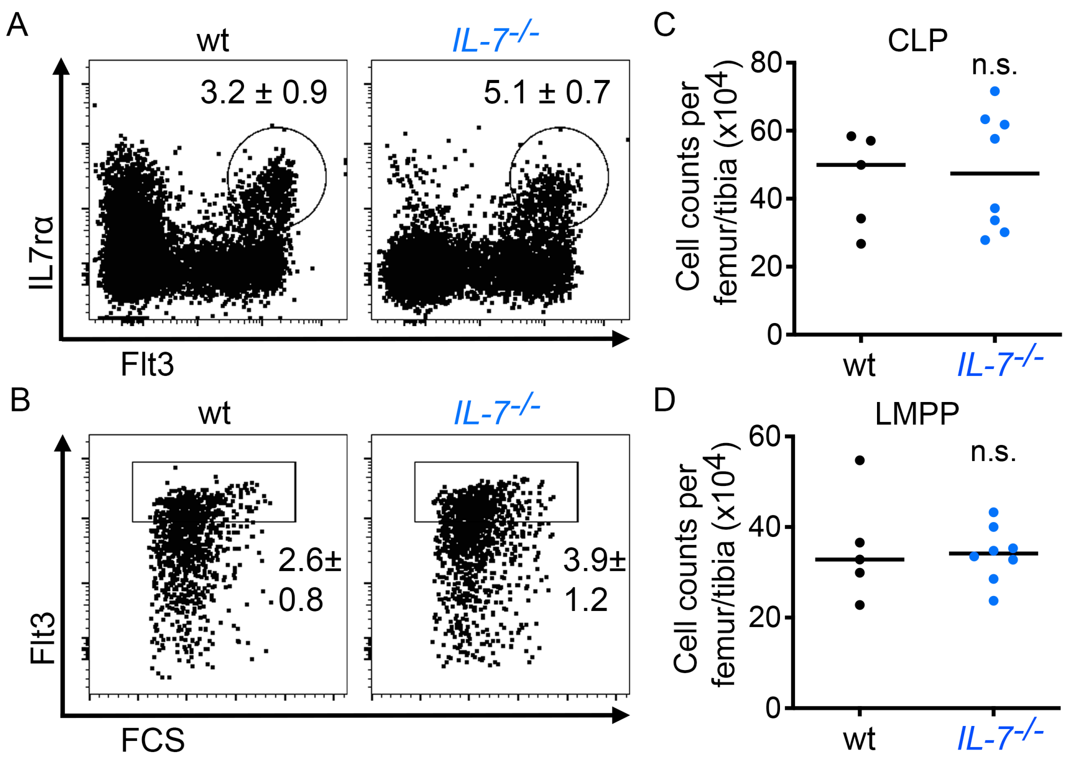


**Figure S1. IL-7-deficient mice have normal numbers of lymphoid-biased progenitors. A)** Representative plots of wild type and IL-7-/- bone marrow cells gated as lineage- showing IL-7rα and Flt3. Gate indicates CLP and the numbers indicate the frequency (mean ± SD) in lineage-negative. **B)** Flt3 expression in wild type and IL-7-/- bone marrow cells gated as LSK (Lineage-Kit+Sca1+). Gate indicates LMPP and the numbers correspond to the frequency (mean ± SD) in lineage-negative. **C-D)** Cellularity of CLPs (C) and LMPPs (D) in the wild type (black) and IL-7-/- (blue) bone marrows. Progenitor counts refer to total cellularity in bone marrow from one femur plus tibia. Mice analyzed were 6 weeks old. Statistical significance was calculated with Mann Whitney test: n.s.p>0.05. Related to Figure 1.

### Figure S2

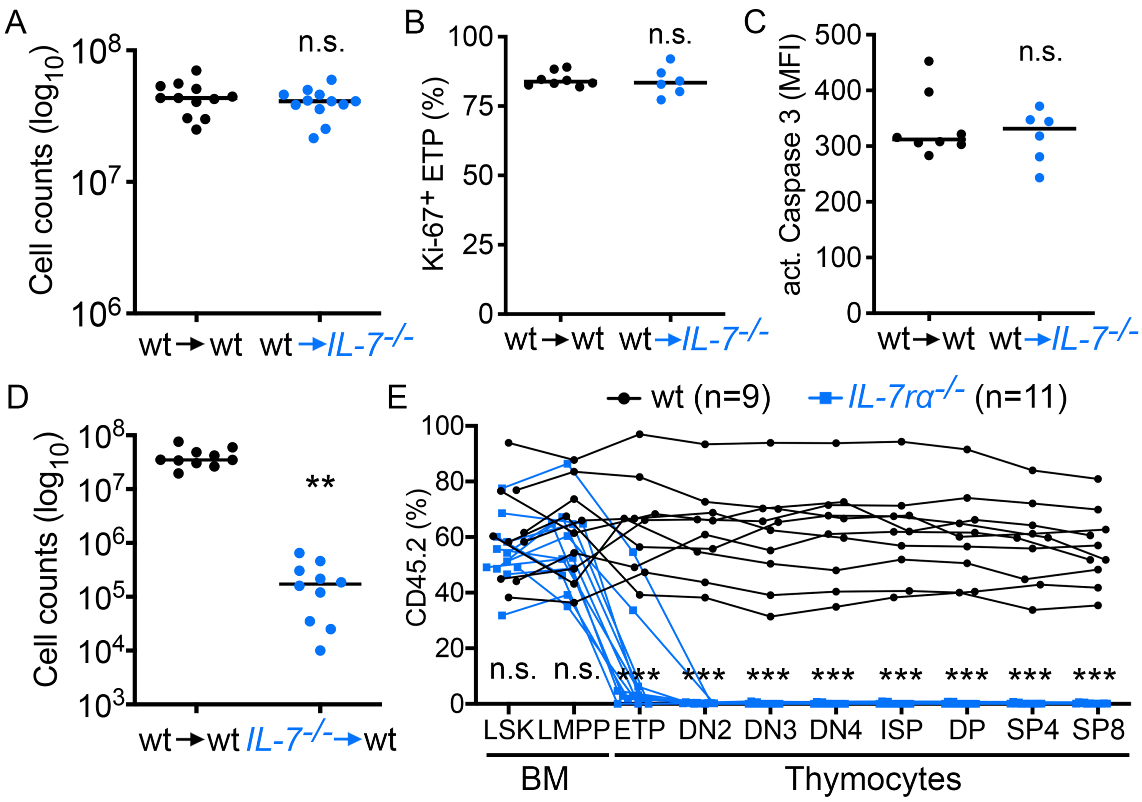


**Figure S2. IL-7r deficiency causes a cell-autonomous defect in ETPs. A)** Total cellularity of wild type thymi transplanted into wild type (black) and IL-7-/- (blue) mice, described in Figure 1D. **B)** Percentage of Ki-67+ ETPs in wild type thymi grafted into wild type (black) or IL-7-/- (blue) mice. **C)** Mean fluorescence intensity of active caspase 3 in ETPs from thymi grafted into wild type (black) or IL-7-/- (blue) mice. **D)** Total cellularity of the wild type (black) and IL-7-/- (blue) thymi grafted into wild type mice, described in Figure 1G. Each symbol represents one mouse (C) or one thymus graft (D-G) and the lines indicate the medians. **E)** Quantification of the frequency of CD45.2+ cells in the chimeras described in Figure 1J (wild type in black and IL-7r -/- in blue) in each population indicated. Competitor CD45.1+ were wild type in both groups and correspond to the percentage missing. Each set of circles connected by the lines corresponds to one mouse. Statistical significance was calculated with Mann Whitney (A-C, E) and paired T (D) tests: ***$p \leq 0.001$, **$p \leq 0.01$, n.s.$p > 0.05$. Related to Figure 2.

### Figure S3

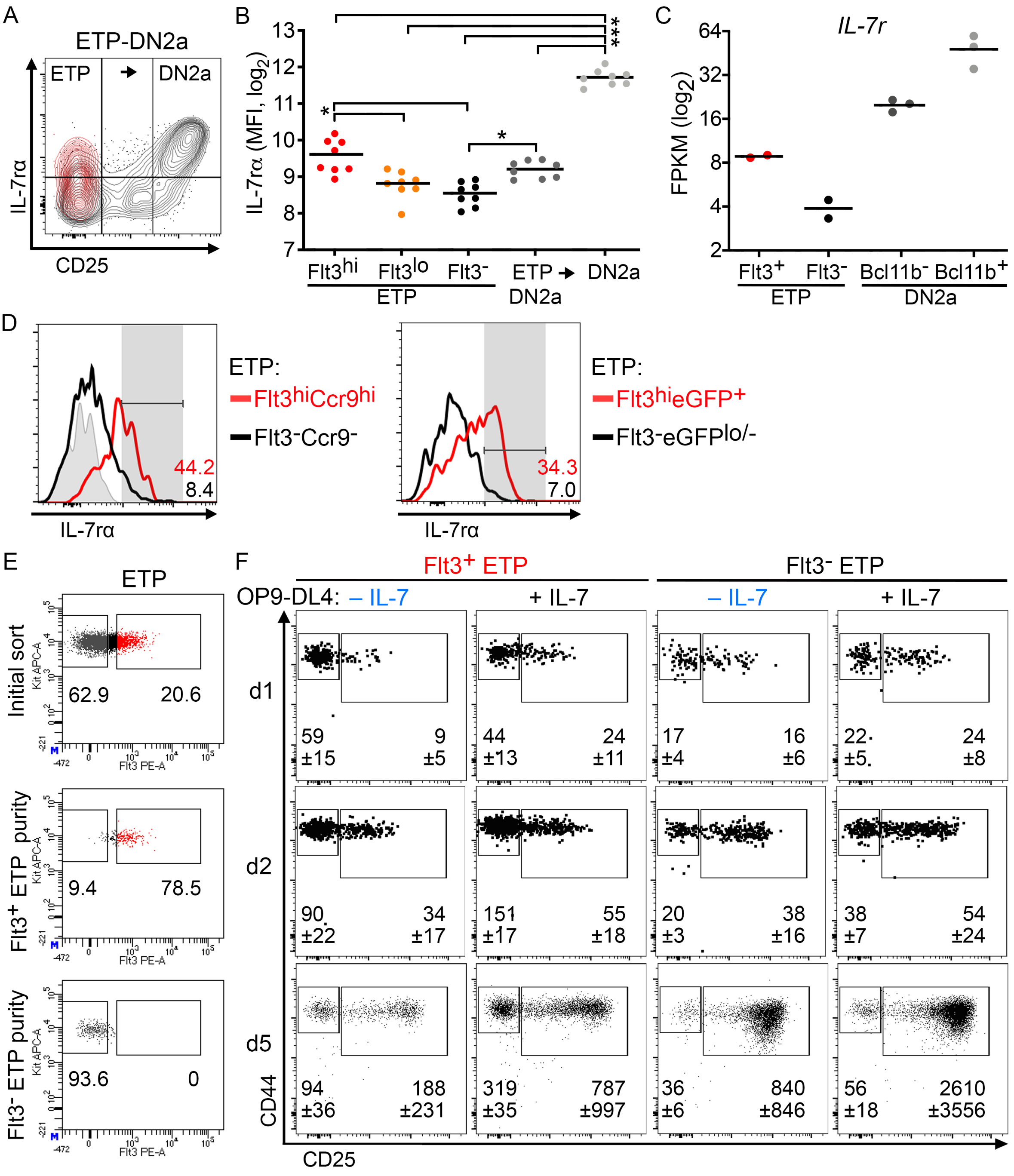


Figure S3. Dynamics of IL-7r expression in early T lymphocyte development. A-B) Thymi from 3-to-5 weeks-old wild type mice were analyzed. A) Contour plots for IL-7ra and CD25 in thymocytes pre-gated in ETP to DN2a (CD4-CD8-Lineage-CD44hiKithi). Flt3hi ETPs are overlaid in red. B) Quantification of the mean fluorescence intensity of IL-7rα in Flt3hi, Flt3lo and Flt3- ETPs, cells transitioning from ETP to DN2a (ETP→DN2a) and DN2a thymocytes. Data are from three independent experiments and each symbol per population represents one thymus. Lines indicate the means and statistical significance was calculated with RM one-way ANOVA with Tukey's multiple comparisons: *p≤0.05, ***p≤0.001. C) Data retrieved from bulk RNAseq produced by another laboratory (Zhou et al., 2019) showing IL-7r FPKMs (fragments per kilobase of transcript per million mapped) in Flt3+ or Flt3- ETPs and in Bcl11b- or Bcl11b+ DN2a. D-E) Flt3+ and Flt3- ETPs were sort-purified and co-cultured with OP9-DLL4 stromal cells with or without IL-7, as indicated. D) IL-7rα expression in the ETP subsets gated in Figure 3D based on co-expression of Flt3 and Ccr9 (left) or eGFP (right). FMO control is shown in grey. E) Gates used for sort of ETPs (pre-gated as CD4-CD8-Lin-CD25-CD44hiKithi) and purities measured after sort for the Flt3+ and Flt3- subsets. F) CD44 and CD25 profiles in live thymocytes (pre-gated as CD45+CD4-CD8-CD3-) developing from co-cultures starting with Flt3+ or Flt3- ETPs with or without IL-7 after 1, 2 or 5 days in culture, as indicated. Gates define ETPs (left) and CD25+ thymocytes (right) and the numbers bellow specify the respective number of cells recovered represented as average ± standard deviation obtained from the replicate wells of one experiment. Plots show the concatenated data from the replicates of the experiment (per condition) and display all the events acquired. Data representative of 3 independent experiments. Related to Figure 4.
